# The temporal organization of corticostriatal communications

**DOI:** 10.1101/2022.07.06.499069

**Authors:** Cole Korponay, Elliot A Stein, Thomas J Ross

## Abstract

The cortex and striatum are linked by hundreds of thousands of structural connections, and the transmission of temporally aligned communications across many of these connections at once is an electrophysiological prerequisite for striatal activation. Despite the importance of communication timing in corticostriatal circuitry, there is little understanding of its system-level organization and properties. To investigate this, we leveraged emerging methods to “temporally unwrap” fMRI data and measure patterns of cortex-striatum node-pair coactivation at frame-wise (i.e., <1 sec) resolution in low head-motion subjects from the Human Connectome Project. First, we identify communities of cortex-striatum node-pairs with preferentially synchronized coactivation patterns. Surprisingly, we find that the map of striatal areas with temporally aligned cortical coactivation patterns does not simply reflect the map of striatal areas with similar cortical connectivity profiles. As a result of the distinct spatial organization of these gradients, striatal nodes connected to similar areas of cortex may nonetheless interact with these cortical areas at different times, and striatal nodes connected to different areas of cortex may nonetheless interact with these areas at similar times. We provide evidence for a possible mechanism driving this divergence: striatal nodes with similar cortical connectivity profiles may have differently timed interactions with cortex if they have different modulatory input profiles (i.e., from the midbrain and thalamus) that differentially gate their responsivity to cortical input. Overall, this blended organization may serve to both increase the repertoire of striatal responses to frontal input and facilitate coordination across functional domains in the temporal dimension. Findings provide a framework to investigate the role of corticostriatal temporal coordination in behavior and disease.

**SIGNIFICANCE STATEMENT:** We provide the first systems-level account of temporal communication patterns between cortex-striatum node-pairs, mapping communities of node-pairs with synchronized communication patterns using an emerging fMRI temporal-unwrapping technique and providing evidence for the mechanism that dictates the organization of these communities. Findings have broad implications for our understanding of the functional architecture of corticostriatal circuits.

## INTRODUCTION

As the model of brain information processing has shifted from one of local functional subunits to distributed networks[1], interest has intensified in examining synchrony between spatially distant neural activity and how this temporal coordination contributes to complex behavioral patterns in both health and disease. To this end, there has been tremendous progress in uncovering pairs and networks of brain nodes with correlated activity[2–4] and the physical tracts that facilitate this synchrony[5, 6]. Such interactions between the frontal cortex and striatum have been afforded particular focus given their centrality in myriad adaptive functions[7, 8] and in neuropsychiatric disorders, including substance use disorder (SUD), obsessive-compulsive disorder (OCD), and attention-deficit hyperactivity disorder (ADHD)[9, 10].

At any given moment, thousands of individual connections transmit communications from the frontal cortex to the striatum. A rich neuroanatomical literature has delineated much of the structural architecture that links nodes in frontal cortex to nodes in striatum[11–13], revealing a topographic organization wherein each cortical region projects preferentially to specific areas of striatum[14]. The transmission of information from frontal cortex to striatum via these projections can be inferred from correlated fMRI BOLD signal activity (i.e., static functional connectivity, FC) between cortex-striatum node-pairs[15, 16]. Though FC is a non-directional measure, and striatal activity also indirectly influences cortical activity via cortico-basal ganglia-thalamo-cortical loops, the FC strength of cortex-striatum node-pairs is strongly correlated to the strength of their direct structural connections. Thus, a topographic map of corticostriatal FC largely mirrors that of the underlying structural connectivity map[16–18]. In aggregate, these corresponding maps of cortex-striatum node-pairs undergird much of the current understanding of interactions between these structures. This framework has engendered an often-implicit assumption that individual cortex-striatum node-pairs are the foundational unit from which behavioral functions emerge in this circuitry.

Yet, morphological and electrophysiological features of frontostriatal circuitry dictate that cortex-striatum node-pairs operate in tandem with one another, via temporal coordination of their communications. Striatal medium spiny neurons (MSNs) have expansive dendritic trees[19] that are subject throughout to a strong inwardly rectifying potassium current that keeps them in a hyperpolarized “down state” at rest[19]. Since individual cortical neurons make only a few synapses on the dendritic tree of any individual MSN[20], even repeated excitatory input from one cortical neuron tends to be insufficient to depolarize an MSN. However, individual MSNs receive afferents from thousands of neurons throughout different areas of cortex[20], and a sufficient number of temporally synchronized communications from these inputs can facilitate depolarization and subsequent firing of action potentials[20–22]. The cortical input to MSNs also tends to be spatially distributed, such that MSNs receive appreciable convergent input from diverse cortical regions[13, 23]. Together, the spatially convergent and temporally coordinated activity of many corticostriatal connections allows the striatum to be an integrator of frontal communications from different functional streams (e.g., motor, cognitive, limbic) and thereby facilitate adaptive, complex behaviors[6]. Overall, communities of cortex-striatum node-pairs (i.e., corticostriatal connections, or ‘edges’ in graph theory) with synchronous patterns of communication appear to represent a crucial component of corticostriatal organization and function.

Yet, the organization and properties of temporal coordination across cortex-striatum node-pairs is poorly understood. Existing structural and functional maps of node-to-node coordination do not provide information about which corticostriatal edges propagate communications that are in sync with one another. High temporal resolution local field potential (LFP) data indicate that the striatum may be spatially differentiable into subregions that have distinct communication patterns with frontal cortex. In particular, evidence exists that oscillation coherence within different cortex-striatum node-pairs occurs at specific preferred frequencies[24], is differentially modulated by the same neurotransmitters[25], and selectively increases for node-pairs actively involved in controlling behavioral output[26]. However, despite the high temporal and spatial resolution of LFP recording techniques, these methods are limited to examining a small subset of the cortical and striatal landscape. As such, a systems-level mapping of the temporal organization of corticostriatal circuitry is lacking.

Emerging methods[27, 28] that allow for time-resolved analysis of “edge” FC interactions, and which can canvass the activity of all of frontal cortex and striatum at millimeter spatial resolution, offer a potential window into this systems-level organization. These methods exploit the finding that much of static FC coherence between two nodes appears to be driven by just a handful of discrete, high-coordination time points – moments that putatively represent bursts of communication between the nodes[29]. Unwrapping the static FC Pearson correlation coefficient between two nodes’ BOLD timeseries (i.e., their *average* level of coactivation across time) can therefore yield an “edge time series” (eTS) that delineates the magnitude of the node-pair’s coactivation at each point in time. This in turn allows for the computation of “edge functional connectivity” (eFC) between any two edges’ eTSs, denoting the synchrony of their coactivation patterns (and putatively, of their communication bursts)[27, 28].

Here, we adapt these methods to elucidate the temporal organization of corticostriatal communications. We first demonstrate the existence of corticostriatal node-pair communities with synchronous coactivation patterns. Surprisingly, we find that the spatial organization of these communities is distinct from the organization of static functional connectivity profiles. Further, we provide evidence that this divergence may stem from spatially diverse modulation of striatal responsivity to cortical input by midbrain and thalamic connections.

## RESULTS

### The Magnitude of Synchrony between Different Corticostriatal Communications

We performed discovery analyses on resting BOLD data from a cohort of 77 unrelated adult subjects from the Human Connectome Project (HCP) dataset[30] curated by Chen and colleagues [31] to meet stringent standards for low head-motion during scanning (see Methods for full cohort details). For each subject we computed the edge time series (eTS, quantifying the magnitude of coactivation between an edge’s two nodes at each time point) for 51,300 corticostriatal edges involving the right striatum (1710 striatal voxels * 30 frontal cortical regions of interest), and for 51,360 corticostriatal edges involving the left striatum (1712 striatal voxels * 30 frontal cortical regions of interest). Frontal cortical regions of interest (ROIs) were derived from the Harvard-Oxford atlas. Subsequently, for each subject, we computed the temporal correlation between every pair of edge time series (edge functional connectivity, eFC), quantifying the magnitude of synchrony between edges’ coactivation patterns (**Figure 1**). The eFC matrix of each subject was averaged together to establish a group mean eFC matrix, upon which subsequent analyses of interest were performed.

**Figure 1.**
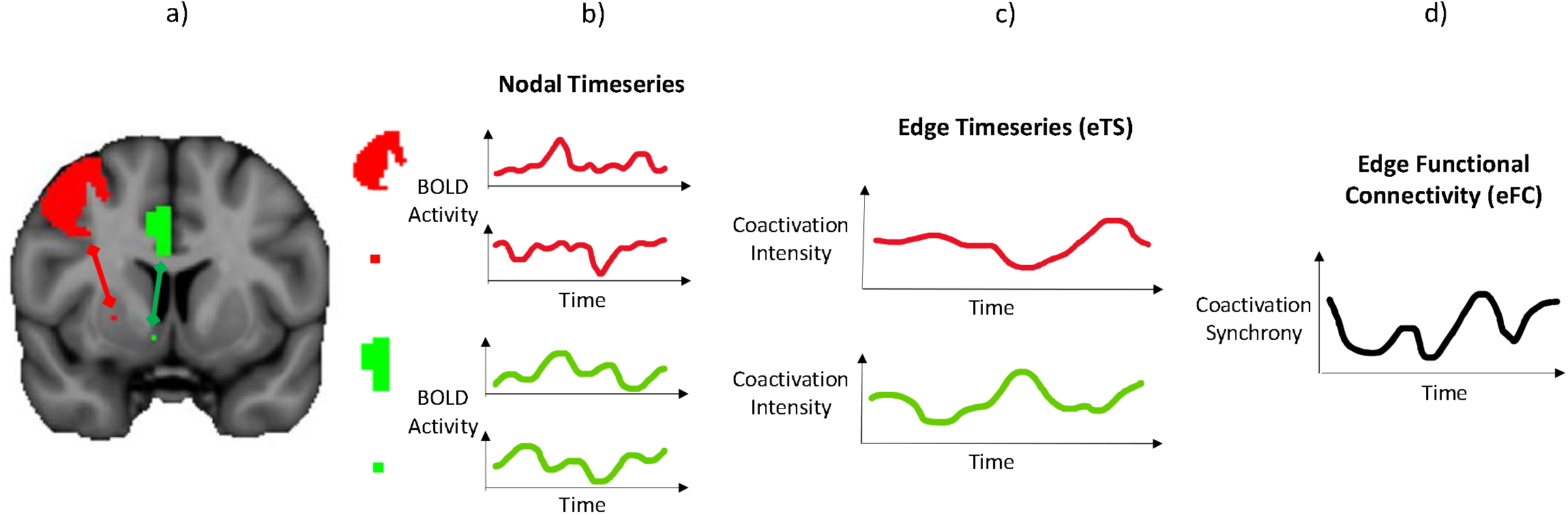
Overview of analysis pipeline. a) Each corticostriatal edge links one cortical node to one striatal node. b) Each node comprising the edge has a “nodal timeseries” quantifying its BOLD activation magnitude across time. c) The correspondence at each time point between the magnitudes of the two nodal timeseries comprising an edge is quantified by the “edge timeseries” (eTS), a putative measure of the communication pattern across the edge. d) Finally, the correlation between any two eTSs is quantified by edge functional connectivity (eFC). We define this as a measure of synchrony between two corticostriatal edges’ communication patterns.

First, we examined the distribution of eFC values across all corticostriatal edge-pairs. We found that 94% of all 1,315,818,540 examined corticostriatal edge-pairs involving the right striatum and 95% of all 1,318,898,310 examined corticostriatal edge-pairs involving the left striatum have very low coactivation synchrony with each other (i.e., edge time series with eFC < 0.2) (**Figure 2**). The preponderance of low-synchrony edge-pairs implies an enormous diversity of node-pair communication patterns. Indeed, on average, a given corticostriatal edge was found to have an eFC > 0.9 with only 2.32 other edges (0.005% of all other edges), and an eFC > 0.5 with only 135.33 other edges (0.264% of all other edges). Overall, eFC values ranged from a low of −0.268 to a high of 0.945. To determine statistically significant eFC values, we established a null distribution of eFC values computed between 2.6 billion random pairs of across-subject eTSs, wherein each eFC value is computed from two eTSs that are statistically independent. p<0.001 corresponded to eFC>0.226 (**Figure S1**).

**Figure 2.**
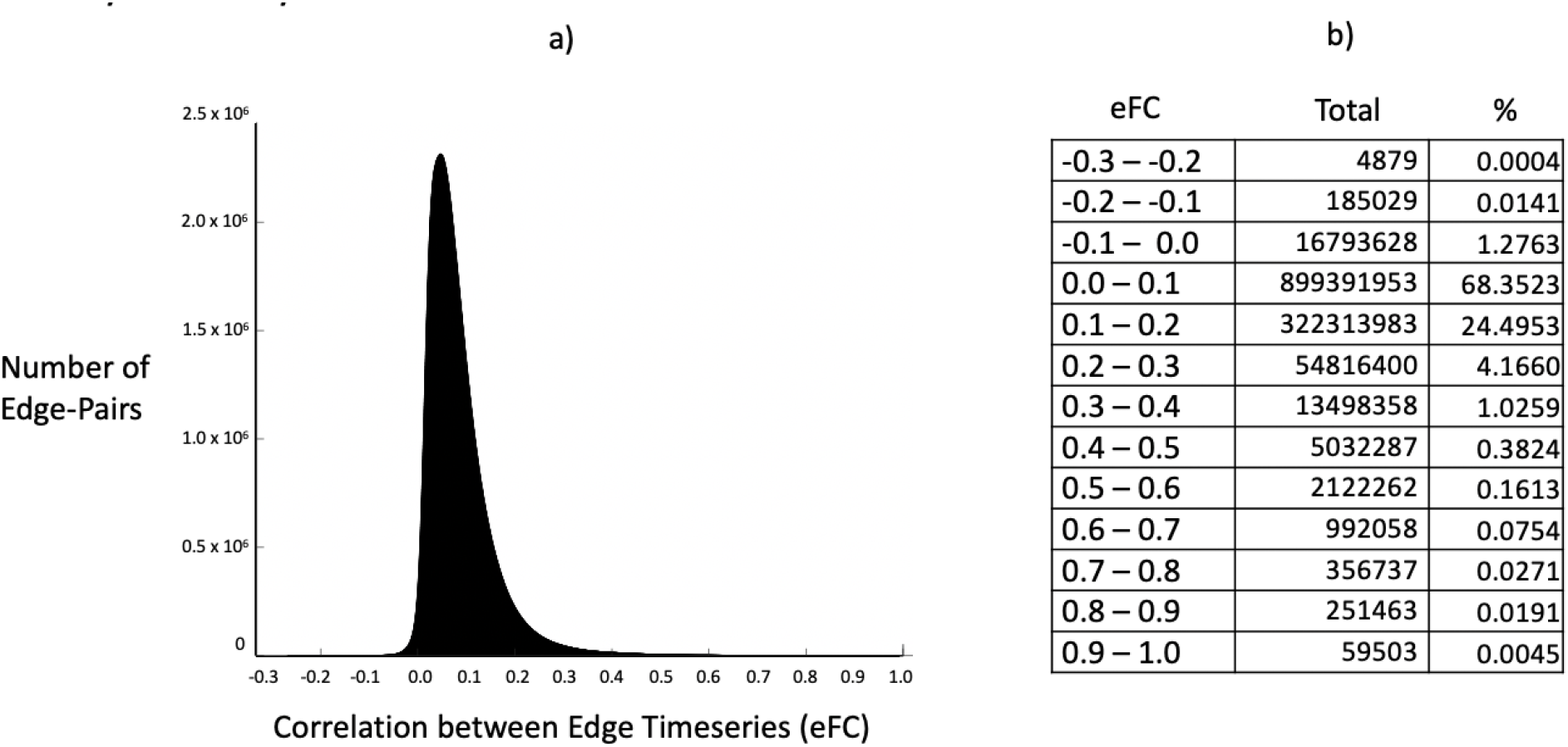
a) Distribution of eFC values across all pairs of corticostriatal edges (right striatum) b) For each range of eFC values, the total number of corticostriatal edge-pairs with an eFC value in that range and the percent of all corticostriatal edge-pairs that total represents.

Next, we examined eFC values amongst different corticostriatal edge-pair “types” (**Figure S2**). We defined five distinct types of corticostriatal edge-pairs. Type 1 edge-pairs are those formed by a single striatal node’s connections to homotopic cortical nodes. Type 2 edge-pairs are those formed via a single striatal node’s connections to non-homotopic cortical nodes. Type 3 edge-pairs are those formed via a single cortical node’s connections to two spatially separated striatal nodes. Type 4 edge-pairs are those formed via two homotopic cortical node’s connections to two spatially separated striatal nodes. Type 5 edge-pairs are those formed via two non-homotopic cortical nodes’ connections to two spatially separated striatal nodes. Amongst these five categories of examined edge-pair types, those formed by a single striatal node’s connections to homotopic cortical nodes (Type 1 edge-pairs) had by far the highest average eFC value (eFC_avg_ = 0.695, p<0.0000001) (**Table 1, Figure S3**), as might be expected given the strong cortico-cortical connections and high level of coordination between homotopic cortical nodes. The next highest average eFC (eFC_avg_ = 0.309, p<0.0001) was amongst edges formed via a single striatal node’s connections to non-homotopic cortical nodes (Type 2 edge-pairs), likely reflecting the electrophysiological requirement of striatal nodes to receive temporally aligned communications from its frontal inputs in order to fire action potentials. In contrast, the magnitude of coactivation synchrony amongst edge-pairs formed by two spatially separated striatal nodes and any combination of cortical nodes (Type 3-5 edge-pairs) was low (eFC_avg_ < 0.175, p<0.01). Such edge-pairs involving either the same cortical node (Type 3, eFC_avg_ = 0.172, p<0.01) or homotopic cortical nodes (Type 4, eFC_avg_ = 0.130, p<0.05) expectedly had coactivation synchrony magnitudes approximately twice as high as those involving two non-homotopic cortical nodes (Type 5, eFC_avg_ = 0.073, p>0.1), whose eTSs were mostly unrelated to each other (**Table 1, Figure S3**).

**Table 1.** Number of edge-pairs involving right striatum in each eFC range by edge-pair type. Bolding indicates each edge-pair type’s most common eFC range. Bottom row presents the mean eFC of each edge-pair type.

|  |  | Edge-Pair Types |  |  |  |  |
| --- | --- | --- | --- | --- | --- | --- |
|  |  | Homotopic<br>Cortical Nodes<br>Same<br>Striatal Node | Non-Homotopic<br>Cortical Nodes<br>Same<br>Striatal Node | Same<br>Cortical Node<br>Spatially Separated<br>Striatal Nodes | Homotopic<br>Cortical Nodes<br>Spatially Separated<br>Striatal Nodes | Non-Homotopic<br>Cortical Nodes<br>Spatially Separated<br>Striatal Nodes |
| Number of<br>Edge-Pairs in<br>Coactivation<br>Synchrony<br>Range | $-0.3 \leq eFC < -0.2$ | 0 | 1676 | 0 | 0 | 135 |
| | $-0.2 \leq eFC < -0.1$ | 0 | 6411 | 0 | 0 | 17332 |
| | $-0.1 \leq eFC < 0.0$ | 0 | 16390 | 1957 | 3586 | 8273959 |
| | $0.0 \leq eFC < 0.1$ | 0 | 86298 | 4479457 | 9378332 | **532055833** |
| | $0.1 \leq eFC < 0.2$ | 0 | **131065** | **13074300** | **12240534** | 145272004 |
| | $0.2 \leq eFC < 0.3$ | 0 | 129108 | 5787553 | 2760382 | 11882810 |
| | $0.3 \leq eFC < 0.4$ | 1660 | 122351 | 1119802 | 426444 | 1583403 |
| | $0.4 \leq eFC < 0.5$ | 2237 | 98727 | 307217 | 112182 | 366713 |
| | $0.5 \leq eFC < 0.6$ | 3791 | 51993 | 111091 | 41544 | 85756 |
| | $0.6 \leq eFC < 0.7$ | 5576 | 47246 | 66083 | 12326 | 18355 |
| | $0.7 \leq eFC < 0.8$ | 3771 | 26913 | 20448 | 4810 | 506 |
| | $0.8 \leq eFC < 0.9$ | **8614** | 22 | 12884 | 355 | 0 |
| | $0.9 \leq eFC \leq 1.0$ | 4 | 0 | 3252 | 0 | 0 |
| Mean eFC |  | 0.695 | 0.309 | 0.172 | 0.130 | 0.073 |

We corroborated the distribution of these observed corticostriatal eFC properties in an independent matched replication sample (n=77) and demonstrated nearly identical correspondence between the eFC matrices of the discovery and replication samples in both the right striatum (r = 0.975, p<0.0001) and left striatum (r = 0.973, p<0.0001) (**Figure S4**). These findings indicate that this temporal organization is highly consistent and reproducible across individuals.

### Coactivation Synchrony Communities: Corticostriatal Connections with Similar Communication Patterns

Next, we sought to identify communities of cortex-striatum node pairs with similar patterns of coactivation across time, as a putative measure of which corticostriatal connections have synchronized communication patterns. To do so, we performed k-means clustering on the group eFC matrix. At all values of k=2 through k=10, k-means clustering identified communities of cortex-striatum node-pairs whose coactivation patterns were significantly (p <0.0001) more synchronized with each other than with node-pairs outside the community. For instance, at k=6, the average eFC between edges within communities was 0.448, 0.410, 0.211, 0.598, 0.458, and 0.501 for Community 1 through Community 6, respectively. Contrarily, the average eFC with edges outside of the community was 0.095, 0.081, −0.047, 0.059, 0.066, and 0.185 for Community 1 through Community 6, respectively. Examination of edge time series at the subject level illustrates the unique temporal coactivation pattern of each community (**Figure 3**), where the average edge time series of each community has a distinct pattern of coactivation magnitude across time.

**Figure 3.**
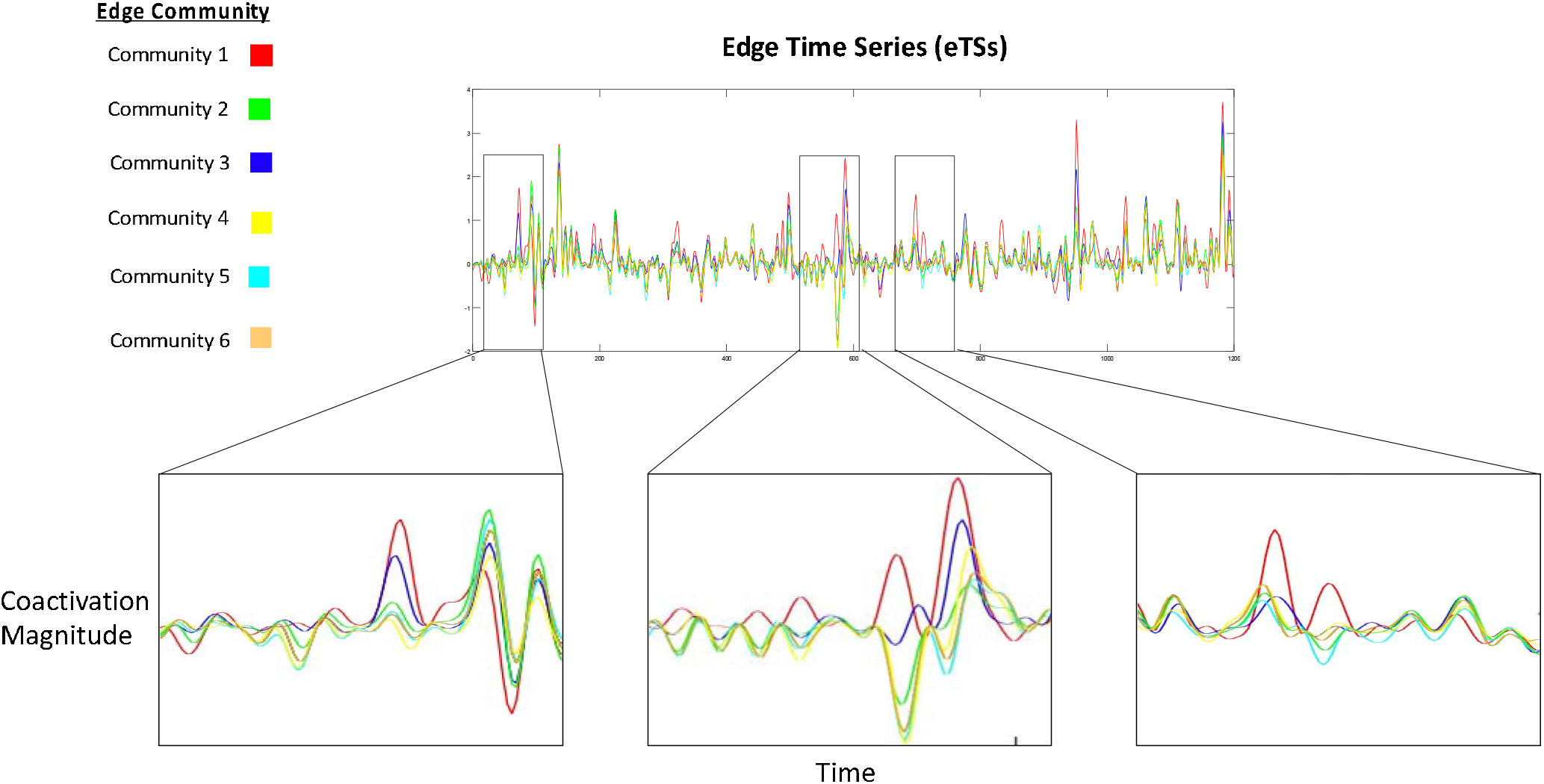
The average edge timeseries (eTS) of edges belonging to each of k=6 coactivation synchrony communities for a representative subject. Edges in each community have a distinct pattern of coactivation magnitude across time.

Each striatal node participates in 30 edges (i.e., one with each of the 30 frontal cortical nodes), and each of these edges belongs to one of *k* coactivation synchrony communities (as determined from the k-means clustering analysis). Thus, each striatal voxel has a multivariate profile of coactivation synchrony community membership (**Figure S5**). In order to facilitate spatial visualization of where corticostriatal connections with similar communication patterns are located, we assigned each striatal voxel to its strongest (i.e., modal) coactivation synchrony community. **Figure 4** illustrates the spatial organization of striatal voxels assigned to their modal coactivation synchrony community, for k=2 through k=10. We found that the number of different strongest communities assigned to striatal voxels reached a maximum at k=6.

**Figure 4.**
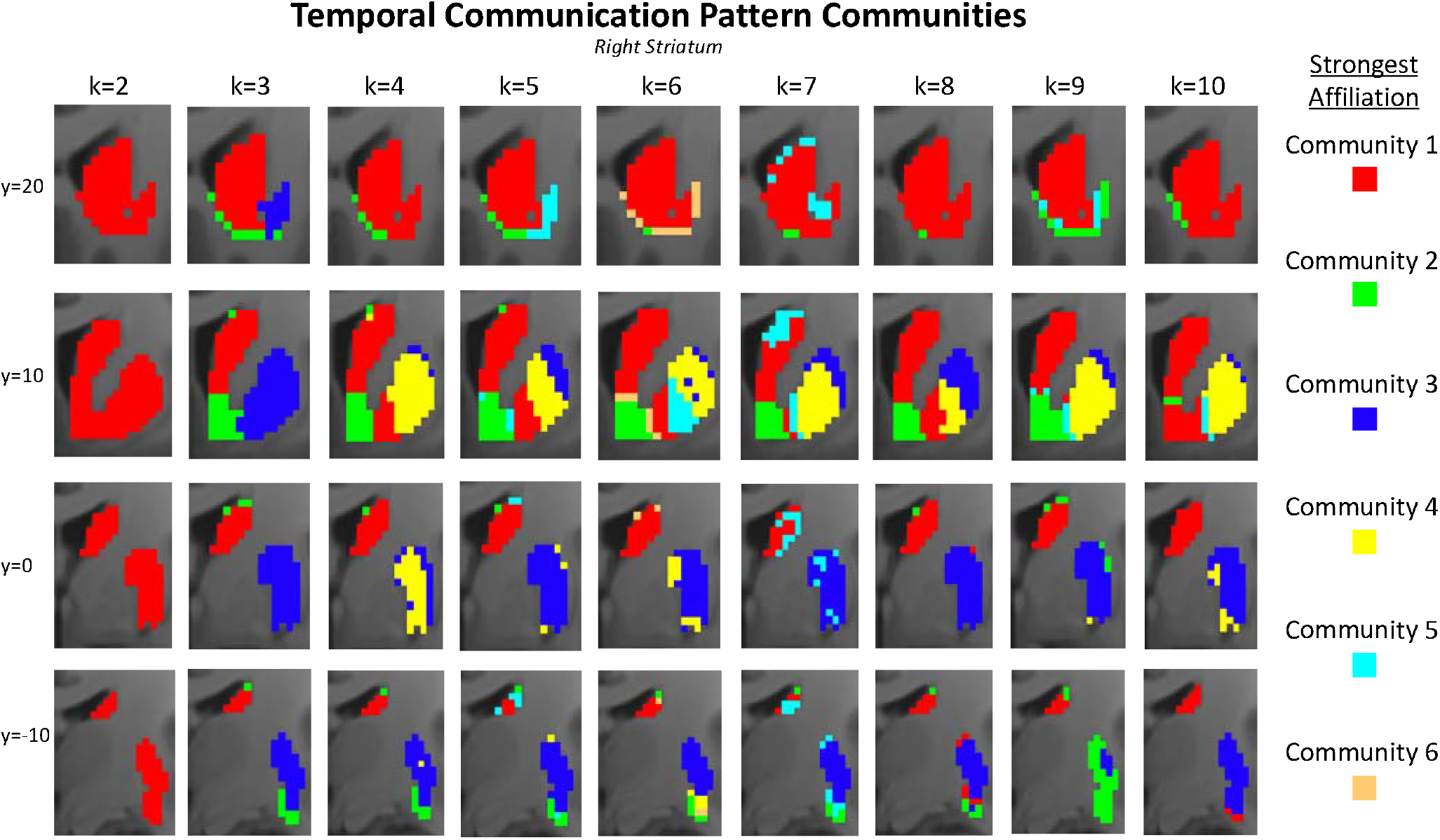
For each integer value of *k*=2 through *k*=10, the modal temporal communication pattern community of each striatal voxel, determined from the community assignments of each voxel’s 30 edges. The number of distinct modal communities reached a maximum of 6 even when *k* exceeded 6.

Common across almost all clustering solutions was 1) a community *(red)* that occupied the rostro-caudal caudate and rostral ventral putamen, 2) a community *(green)* that occupied the nucleus accumbens; and 3) a community *(dark blue)* that occupied the caudal putamen. The putamen contained the greatest diversity of temporal communication pattern communities. In contrast to the caudate and nucleus accumbens, which each contained primarily one temporal communication pattern community even as *k* increased, the putamen consistently subdivided into as many as three distinct temporal communication pattern communities at *k*=4 and above.

### Comparing Temporal Communication Pattern Communities to Connectivity Profile Communities

Next, we sought to examine whether the spatial distribution of striatal temporal communication pattern communities mirrors that of striatal connectivity profile communities or manifests as a distinct organizational gradient. To do so, we used k-means clustering (k=2 through k=10) to cluster striatal voxels into communities with similar static functional connectivity profiles (with the 30 examined ROIs of frontal cortex). We then compared these maps to the temporal communication pattern maps. Importantly, the static functional connectivity values used to establish the connectivity profile communities were computed from the same edge time series data that were used to compute the eFC values that defined the temporal communication communities. We validated that static functional connectivity values computed from edge time series data are precisely equivalent to static functional connectivity values computed through traditional methods (**Figure S6**).

Overall, while connectivity profile and communication pattern communities are strongly aligned in the nucleus accumbens, these gradients have distinct spatial organizations in the caudate and putamen that become increasingly apparent at higher values of *k*. Most notably, a single communication pattern (red) dominates the caudate and rostral ventral putamen, even though these areas consist of diverse connectivity profile communities; vice versa, multiple communication patterns (red, yellow, blue) subdivide areas of the precommissural putamen that consist of single connectivity profile communities (**Figure 5**).

**Figure 5.**
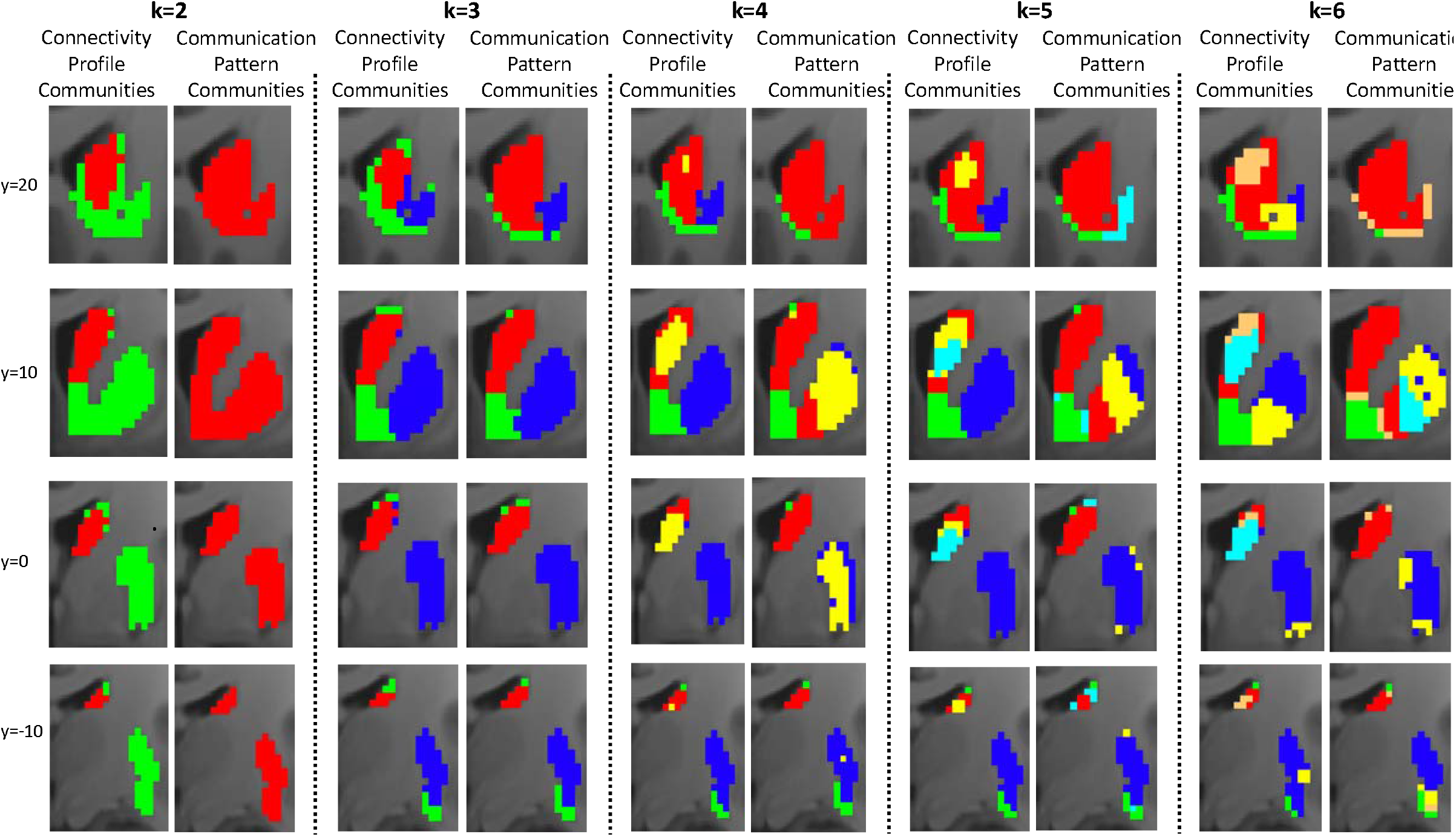
Comparing the spatial organization of frontal cortical connectivity profile communities (left panel) and frontal cortical communication pattern communities (right panel) at different levels of *k*.

This divergence can also be concretely observed at the single subject and single ROI level, wherein striatal voxels with similar static functional connectivity strengths with a frontal cortical ROI can nonetheless have different patterns of communication with it (**Figure S7**). The ability of these parameters to diverge can be understood by considering that the same average value (i.e., a static functional connectivity value) can be generated by multiple different number combination sets (i.e., patterns of temporal coordination magnitudes). This potential for diversity in the relationship between average values and their underlying constituent values appears to manifest in corticostriatal interactions, such that time-averaged connectivity strength and temporal pattern of communication are dissociable parameters of a striatal node’s interaction with a frontal cortical ROI.

These findings beg a further question. How can two striatal nodes that have similar frontal cortical connectivity profiles – and which therefore receive similarly timed communications from similar inputs – nonetheless have different temporal patterns of interaction with these inputs?

### Variance in Modulatory Input Profiles as a Candidate Mechanism for Decoupling Corticostriatal Static Connectivity from Corticostriatal Temporal Dynamics

In addition to frontal input, striatal nodes receive substantial input from other areas of the striatum, the dopaminergic midbrain, and the thalamus. Moreover, each of those non-cortical input sources have been found to modulate, or “gate”, the responsivity of striatal nodes to cortical input[32–36]. Thus, theoretically, two striatal nodes receiving the same inputs from cortex could have distinct coactivation patterns with these areas if they receive different profiles of modulatory input, which would gate their responsivity to the cortical input in separable manners. For the data to support such a mechanism, we should expect to see signatures of the frontal communication pattern community organization (which are not present in the frontal connectivity profile community organization) in the connectivity profile community organization of one or more of the non-frontal modulatory inputs.

Indeed, features of the communication pattern map (**Figure 6e**) that are not found in the frontal cortical static FC map (**Figure 6d**) are identifiable in one or more of the modulatory static FC maps (**Figure 6a-c**). For instance, the communication pattern community (red, **Figure 6e**) that spans both the rostral caudate and rostral ventral putamen is identifiable in both the midbrain-striatal (red, **Figure 6b**) and thalamic-striatal (red, **Figure 6c**) static FC maps. In these regions, striatal nodes with different cortical connectivity profiles (blue, yellow, red, copper **Figure 6d**) nonetheless have similar midbrain and thalamic connectivity profiles (red, **Figure 6b-c**). The alignment of this midbrain and thalamic connectivity profile community (red, **Figure 6b-c**) with the frontal communication pattern community (red, **Figure 6e**) suggests that these modulatory inputs may play a role in shaping this communication pattern and its spatial extent. As another example, the partition of the rostral ventral putamen into more communication pattern communities (**Figure 6e**) than frontal connectivity profile communities (**Figure 6d**) at each level of *k* is also reflected in the midbrain and thalamic connectivity profile maps (**Figure 6b-c**). Here, a region characterized by similar frontal connectivity nonetheless has diverse midbrain and thalamic connectivity. Again, the alignment of these midbrain and thalamic connectivity profile communities with frontal communication pattern communities is suggestive of their role in shaping them. Notably, the modulatory connectivity profiles of the nucleus accumbens (green, **Figure 6a-c**) are unique in all being spatially aligned with the region’s frontal connectivity profile (green, **Figure 6d**). This might explain why it is also the only region of the striatum to have spatial alignment in its frontal connectivity profile and frontal communication pattern.

**Figure 6.**
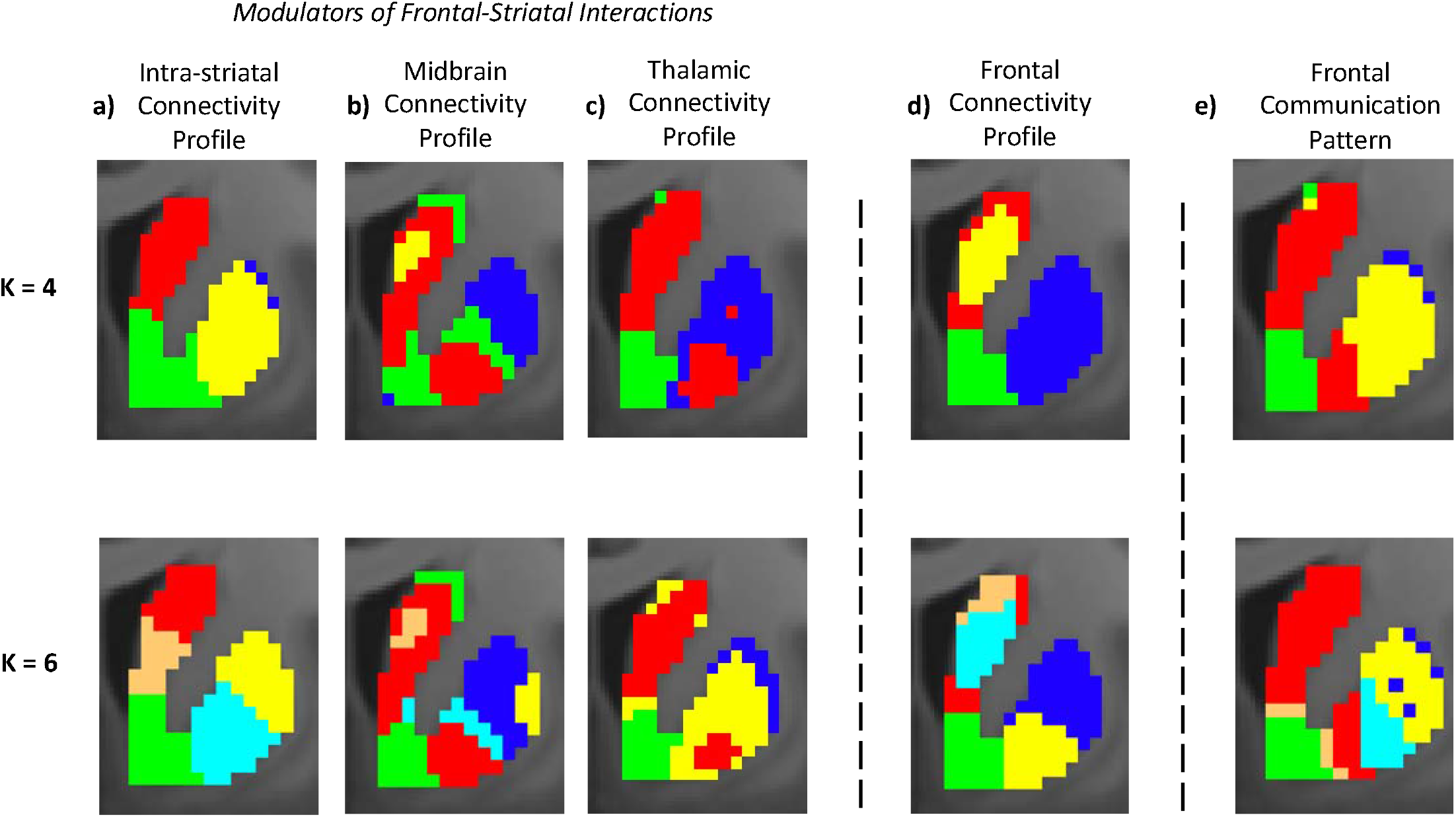
Visualizing the spatial alignment between striatal nodes with similar a) intra-striatal static FC profiles, b) midbrain-striatal static FC profiles, c) thalamo-striatal static FC profiles, d) frontal cortical static FC profiles, and e) frontal cortical communication patterns.

Collectively, these data suggest that two striatal nodes with similar frontal cortical connectivity profiles – receiving similarly timed communications from similar frontal inputs – may nonetheless have different frontal interaction patterns if they have different modulatory input profiles that differentially gate their responsivity to cortical input.

### The Synchrony of Individual Striatal Nodes’ Communications

Lastly, we investigated spatial variance in the coherence each striatal voxel’s set of frontal cortical communications. We defined “synchrony” as the average eFC amongst all paired combinations of a given striatal voxel’s 30 frontostriatal edge time series. This measure provides a putative gauge of how aligned a striatal node’s frontal cortical communications are with each other. We found that a dorsomedial to ventrolateral diagonal strip through the central striatum displayed elevated levels of communication synchrony compared to the rest of the striatum (**Figure S8**). This may indicate that these areas have a relatively higher number of cortical inputs that frequently provide temporally coordinated input.

## DISCUSSION

The Hebbian principle holds that brain nodes that are active at the same time are likely involved in a common functional process[39]. In an edge-centric interpretation of this principle, connections that transmit communications at the same time are also likely involved in a common functional process. This notion comes to bear concretely in the striatum, where individual MSNs must receive temporally aligned communications from many of their afferents as a prerequisite for firing action potentials[20]. Yet, despite extensive knowledge of the path and strength of corticostriatal communications[6, 17], there has been little understanding of which communications tend to be transmitted concurrently with one another. Put another way, while the literature has elucidated a tremendous amount about coordination within cortex-striatum node-pairs, comparatively little is known about coordination between them.

Here, we computed moment-by-moment (albeit, at a resolution of 720 msec) coactivation magnitudes in cortex-striatum node-pairs (edges) – between which communications are transmitted across well-characterized monosynaptic connections – as a proxy for temporal patterns of interaction intensity. We found that there is a tremendous diversity of corticostriatal communication patterns, as >94% of corticostriatal edge-pairs have coactivation patterns with low synchrony (eFC < 0.2). Nonetheless, corticostriatal node-pairs can be robustly and reproducibly clustered into a small set of communities with similar communication patterns. The spatial organization of these communication pattern communities is not simply a reflection of the striatum’s connectivity profile communities. Nodes throughout the caudate and rostral ventral putamen that have different frontal connectivity profiles nonetheless share a similar frontal communication pattern. Moreover, nodes throughout the precommissural putamen that have similar frontal connectivity profiles nonetheless have different frontal communication pattern. Indeed, only in the ventromedial striatum do striatal nodes with similar frontal connectivity profiles also consistently have similar frontal communication patterns. Collectively, the distinct spatial organization of these gradients produces a structure in which 1) striatal nodes connected to similar areas of cortex may nonetheless communicate with these areas at different times and 2) striatal nodes connected to different areas of cortex may nonetheless communicate with these areas at similar times. This spatiotemporal “blending” may serve to both increase the repertoire of striatal responses to frontal input within functional domains and facilitate coordination across functional domains in the temporal dimension. This type of “blended” circuit organization, wherein gradients of distinct modalities are oriented distinctly from one another to create a matrix of diverse computational units, has been observed in the visual cortex[40]. Here, orientation columns are organized orthogonally to ocular dominance columns, with different color “blobs” embedded throughout the matrix, to create distinct functional units to process different combinations of orientation, eye and color input[41]. Moreover, this matrix of computational units is highly plastic, both in the context of development and in response to changes in visual input[42]. This exemplar suggests that the spatiotemporal blending observed here in the striatum may establish a previously unrecognized matrix of computational units that warrants future examination in the context of behavior and development.

How might striatal nodes that receive similarly timed communications from similar frontal areas nonetheless have different patterns of coactivation with these frontal areas? We hypothesized that other striatal afferents known to modulate or “gate” striatal responsivity to frontal cortical input might play a role in this phenomenon. This hypothesis was based on several neurobiological observations. First, within the striatum, MSNs receive an intricate set of modulatory input both from other MSNs via axon collaterals and from a diverse set of striatal interneurons [32–34]. It has been posited that lateral inhibition between MSNs is foundational in regulating MSN excitability to frontal input and shaping behaviorally relevant ensembles of MSNs[34]. The striatum also receives afferents from the bilateral dopaminergic midbrain and thalamus that are known to modulate MSN excitability to frontal cortical input[35, 36]. For instance, thalamic glutamatergic input to striatal cholinergic interneurons is thought to gate the influence of cortical input on striatal MSNs[36], and dopaminergic projections from the substantia nigra and ventral tegmental area are known to directly modulate MSN excitability via interactions with MSN D1 and D2 receptors[35]. Thus, theoretically, if striatal MSNs with similar connectivity profiles with frontal cortex have different profiles of intra-striatal, midbrain-striatal and thalamo-striatal connectivity, these MSNs could have different magnitudes of responsivity to the same frontal input at a given moment in time.

We provide evidence of such a model in our data, as spatial features of the frontal cortical communication pattern map that are not found in the frontal cortical static FC map are identifiable in the striatum’s static FC maps with the midbrain and thalamus. The alignment of the midbrain and thalamic connectivity profile communities with the frontal communication pattern community suggests that these modulatory inputs may play a role in shaping the frontal communication pattern and its spatial extent. Overall, these data suggest that two striatal nodes with similar frontal cortical connectivity profiles – receiving similarly timed communications from similar frontal inputs – may nonetheless have different frontal interaction patterns if they have different modulatory input profiles that differentially gate their responsivity to cortical input. However, it should be noted that our examination of these processes is limited by the temporal (720 msec) and spatial (2 mm^3^) resolution of fMRI. Functional units of both time and space occur at smaller scales in the striatum[43], and thus follow-up work using electrophysiological techniques is warranted.

Overall, we provide evidence for a systems-level temporal organization of corticostriatal circuitry. Findings and methods provide a framework for using fMRI to investigate how different corticostriatal connections coordinate – via temporal synchrony of their communications – to produce complex behaviors and how these may be altered in various neuropsychiatric diseases.

## METHODS

Discovery dataset analyses were performed on rsfMRI data from a cohort of 77 unrelated adult subjects (age: 29.09 ± 3.86; 36.4% male; all right-handed) from the Human Connectome Project (HCP) dataset[30] curated by Chen and colleagues [31] to meet the following stringent standards for low head-motion during scanning: (1) range of head motion in any translational direction less than 1 mm and (2) average scan-to-scan head motion less than 0.2 mm. An age-, gender- and handedness-matched n=77 replication sample (age: 29.08 ± 3.62; 33.8% male; all right-handed) was also drawn from the HCP dataset. Sample matching was performed using nearest neighbor matching via the R program MatchIt (https://cran.r-project.org/web/packages/MatchIt/MatchIt.pdf). Briefly, the program computed a distance between each discovery sample subject and the remaining HCP subjects, and, one-by-one, selected a match for each discovery sample subject. The HCP study was approved by the Washington University Institutional Review Board and informed consent was obtained from all subjects.

### fMRI Data Acquisition

HCP neuroimaging data were acquired with a standard 32-channel head coil on a Siemens 3T Skyra modified to achieve a maximum gradient strength of 100 mT/m [30, 44, 45]. Gradient-echo EPI images were acquired with the following parameters: TR = 720 ms, TE = 33.1 ms, flip angle = 52º, FOV = 280 × 180 mm, Matrix = 140 × 90, Echo spacing = 0.58 ms, BW = 2,290 Hz/Px. Slice thickness was set to 2.0 mm, 72 slices, 2.0 mm isotropic voxels, with a multiband acceleration factor of 8. Resting state data used in the present analyses were acquired from runs of approximately 14.4 min each (REST1). Participants were instructed to lie still with their eyes open and fixated on a bright crosshair on a dark background.

### Preprocessing

We began with HCP minimally preprocessed resting-state data [46]. Briefly, this preprocessing pipeline removes spatial distortions, realigns volumes to compensate for subject motion, registers the echo planar functional data to the structural data, reduces the bias field, normalizes the 4D image to a global mean, and masks the data with a final FreeSurfer-generated brain mask [46] We performed further preprocessing steps on each subject’s resting-state data including spatial blurring with a 6-mm full-width half-maximum Gaussian kernel and temporal filtering (0.01<f <0.1 Hz). To further minimize effects of subject head-motion, volumes were censored for framewise motion displacement (i.e., volume to volume movement) >0.5 mm [47, 48]. Finally, the time series of each voxel was confound regressed in a GLM with 17 regressors of no interest: six motion parameters (three translations and three rotations) obtained from the rigid-body alignment of EPI volumes and their six temporal derivatives; the mean time series extracted from white matter; the mean times series extracted from CSF; and a second-order polynomial to model baseline signal and slow drift.

To extract the frontal cortical and striatal timeseries of interest, we performed the following steps. First, we parcellated each hemisphere of the frontal cortex into 15 regions of interest (ROIs) using the Harvard-Oxford atlas (http://www.fmrib.ox.ac.uk/fsl/): supplementary motor cortex, superior frontal gyrus, subcallosal cortex, precentral gyrus, paracingulate gyrus, middle frontal gyrus, insular cortex, inferior frontal gyrus pars opercularis, inferior frontal gyrus pars triangularis, frontal pole, frontal orbital cortex, frontal operculum cortex, frontal medial cortex, central opercular cortex, and anterior cingulate cortex. For each subject, a mean timeseries was computed from the voxels comprising each of the 30 ROIs. Sample-specific right and left striatum masks were created by averaging the union of the caudate, putamen, and nucleus accumbens FreeSurfer parcellations of each subject. This resulted in right and left striatum masks of 1710 and 1712 voxels, respectively; a timeseries was extracted from each voxel. As such, each subject had 1710 right striatal timeseries, 1712 left striatal timeseries, and 30 frontal cortical timeseries. Each of these timeseries was Z-scored. Since we sought to investigate corticostriatal connections and not striatal-striatal connections, analyses were performed separately for the right and left striatum.

### Analytic Strategy

#### Building the Frontostriatal eFC Matrix

To compute corticostriatal edge time series (eTS) and edge functional connectivity (eFC) matrices[27] for each subject, we performed the following steps (see **Figure 1**). For each of the 1710*30=51,300 unique cortical node-striatal node pairs (edges) in the right striatum, we computed the element-wise product of the pair’s Z-scored timeseries. The resulting 51,300 eTSs each encoded the magnitude by which activity of the two nodes comprising the edge was synchronized at each moment in time (i.e., at each TR). Increasingly positive values in the eTS represent moments where activity in the two nodes is more synchronized (i.e., where activity is either high or low at both nodes). In contrast, increasingly negative values in the eTS represent moments where activity in the two nodes is less synchronized (i.e., where activity at one node is high while activity in the other node is low). Lastly, the scalar product of each pair of eTSs was taken to compute a 51,300 × 51,300 eFC matrix. Each value in the eFC matrix represents the correlation between the synchrony magnitudes of two eTSs across time. This provides an index of how similar the temporal pattern of nodal activity synchrony is for two distinct edges. The eFC matrix of each subject was averaged together to establish a group mean eFC matrix, upon which subsequent analyses of interest were performed. An analogous process was undertaken for the left striatum.

#### Analyzing the eFC Matrix

To cluster edges with similar eTSs into “coactivation synchrony communities”, we used a standard *k*-means algorithm with squared Euclidean distance and 1,000 iterations at each value of *k*=2 through *k*=10. To quantify the diversity of each striatal voxel’s coactivation synchrony community profile, we computed each profile’s entropy, which indexes the uniformity of its distribution of values:

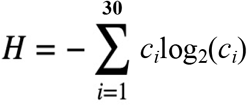

where *c_i_* is the community number the voxel’s *i*-th edge[38]. For each voxel we divided this measure by log_2_30 to normalize it and bind it to the interval [0,1][49]. Higher values of entropy indicate voxels whose edges belong to many different communities. Conversely, lower values of entropy represent voxels whose edges belong primarily to the same community.

#### Defining Edge-Pair Types

To conduct descriptive analyses of the properties of coactivation synchrony across different kinds of corticostriatal connections, we categorized edge-pairs into five “types” (**Supplementary Figure 1**). To define “spatially separated” striatal nodes, we parcellated the striatum into six non-overlapping sub-territories: rostral and caudal caudate, rostral and caudal putamen, and rostral and caudal nucleus accumbens. The Oxford-GSK-Imanova Structural–Anatomical Striatal Atlas[50] (https://identifiers.org/neurovault.image:406338) was used to guide delineation of caudate, putamen and nucleus accumbens; the anterior commissure was used as an anatomical landmark to delineate rostral versus caudal subdivisions for the caudate and putamen; subdivisions of the nucleus accumbens were determined using the first and second half of coronal slices where the structure appeared. Two striatal nodes situated in two different sub-territories were considered to be “spatially separated”; pairs of striatal nodes where one was located on or adjacent to the border of two sub-territories were not considered as spatially separated. Edges formed between cortical nodes and adjacent striatal nodes were not considered in this “types” framework, given the highly correlated time series of adjacent striatal voxels.

#### Examining Intra-Striatal, Fronto-Striatal, Midbrain-Striatal, and Thalamo-Striatal Static Functional Connectivity

The correlation between the BOLD timeseries of each striatal voxel and the BOLD timeseries of each of the following were computed: 1) all remaining voxels in the striatum, 2) the 30 frontal cortical ROIs, 3) all voxels in the bilateral midbrain, and 4) all voxels in the bilateral thalamus. Voxel-wise time-series from the midbrain were extracted from masks of the bilateral[51] substantia nigra and ventral tegmental area as defined by Murty et al [52, 53] (https://neurovault.org/images/46839/). Voxel-wise time-series from the bilateral thalamus were extracted from the mask defined by Krauth et al [54, 55] (https://neurovault.org/images/111302/). Thus, each striatal voxel was assigned four independent static connectivity profiles – one with each of the four structures (striatum, frontal cortex, midbrain, and thalamus). K-means clustering was performed separately for striatal connectivity profiles with each of the four structures, to determine the distinct spatial distribution of striatal connectivity profiles with each structure.

## DATA AVAILABILITY

All imaging data come from the Human Connectome Project’s publicly available, open access repository at https://db.humanconnectome.org/app/template/Login.vm, which can be accessed after signing a data use agreement.

## CODE AVAILABILITY

Code to apply “temporal unwrapping”/edge functional connectivity and its related derivatives to corticostriatal circuits or other circuits/networks of interest has been made available at https://github.com/ckorponay/Temporal_Organization_of_Communications.

## Acknowledgements

Data were provided by the Human Connectome Project, WU-Minn Consortium (Principal Investigators: David Van Essen and Kamil Ugurbil; 1U54MH091657) funded by the 16 NIH Institutes and Centers that support the NIH Blueprint for Neuroscience Research, and by the McDonnell Center for Systems Neuroscience at Washington University. Funding for this work was provided by the National Institute on Drug Abuse (1F32DA048580-01A1 to C.K.) and the Intramural Research Program of the National Institute on Drug Abuse (to T.R. and E.A.S.).

