## Supplemental Material for "The temporal organization of corticostriatal communications"

**SUPPLEMENTARY MATERIALS**

Figure S1.

**Null eFC Distribution**

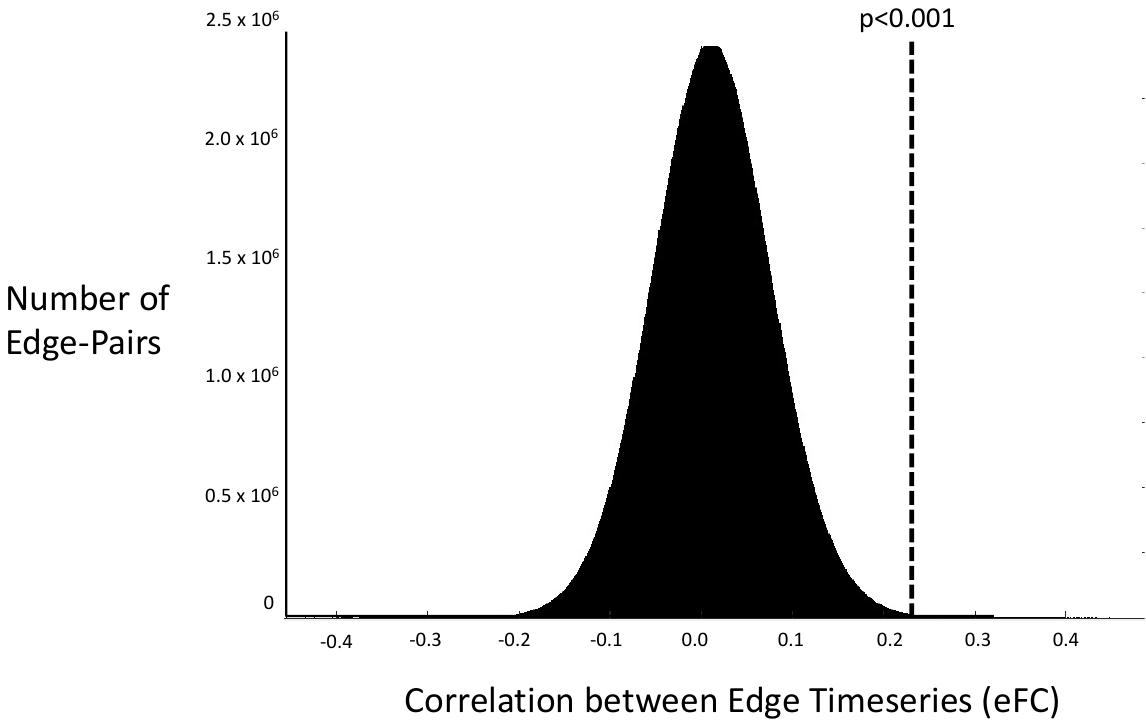

**Figure S1.** The null distribution of 2.6 billion eFC values computed from randomly permuted across-subject (i.e., independent) edge time series. Demonstrates the likelihood of observing each eFC value by random chance, and provides a p<0.001 significance threshold of eFC > 0.226.

Figure S2.

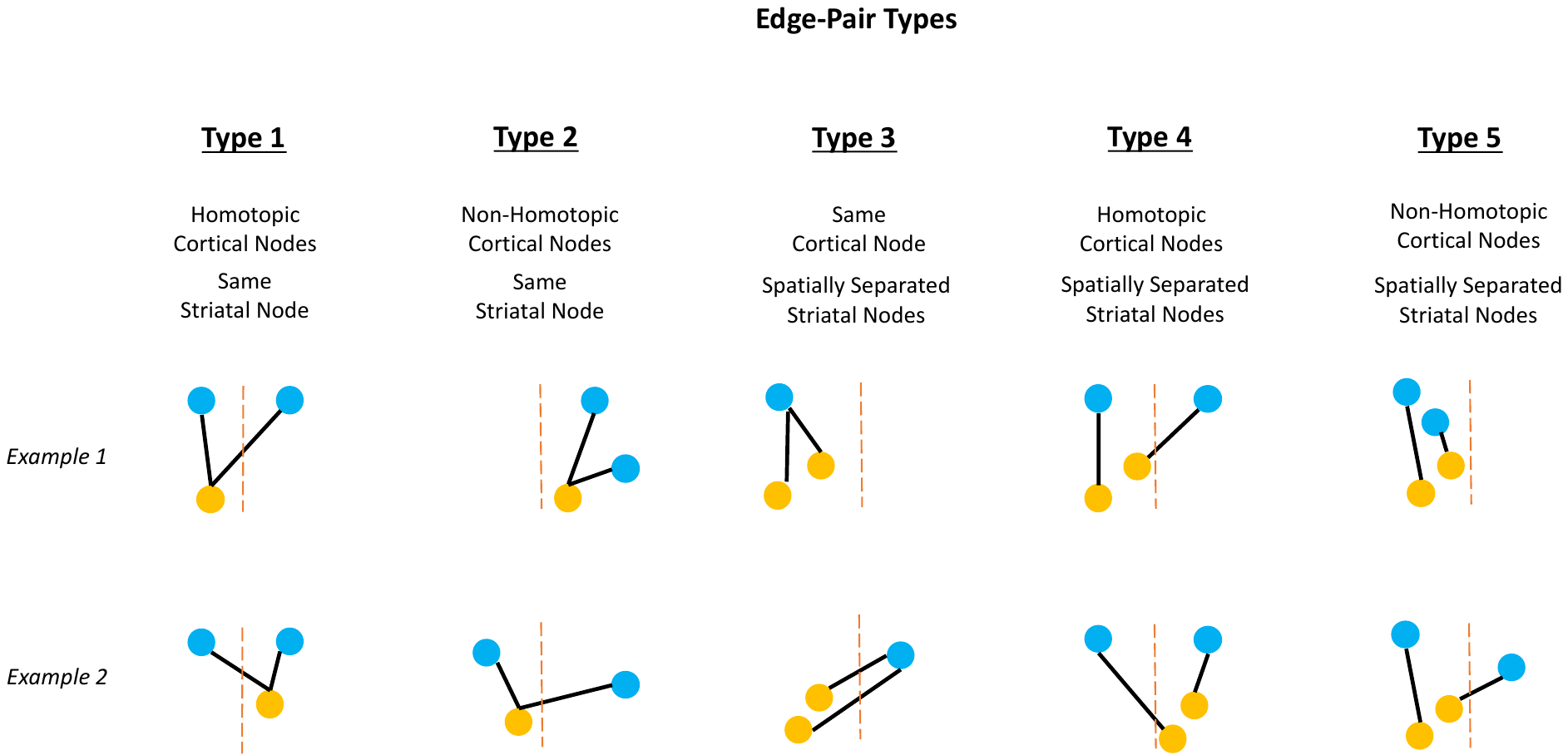

**Figure S2**. The five distinct types of corticostriatal edge-pairs examined; two examples of each. The dashed line indicates the interhemispheric fissure. Edge-pairs involving striatal nodes in different hemispheres were not considered for this analysis.

Figure S3.

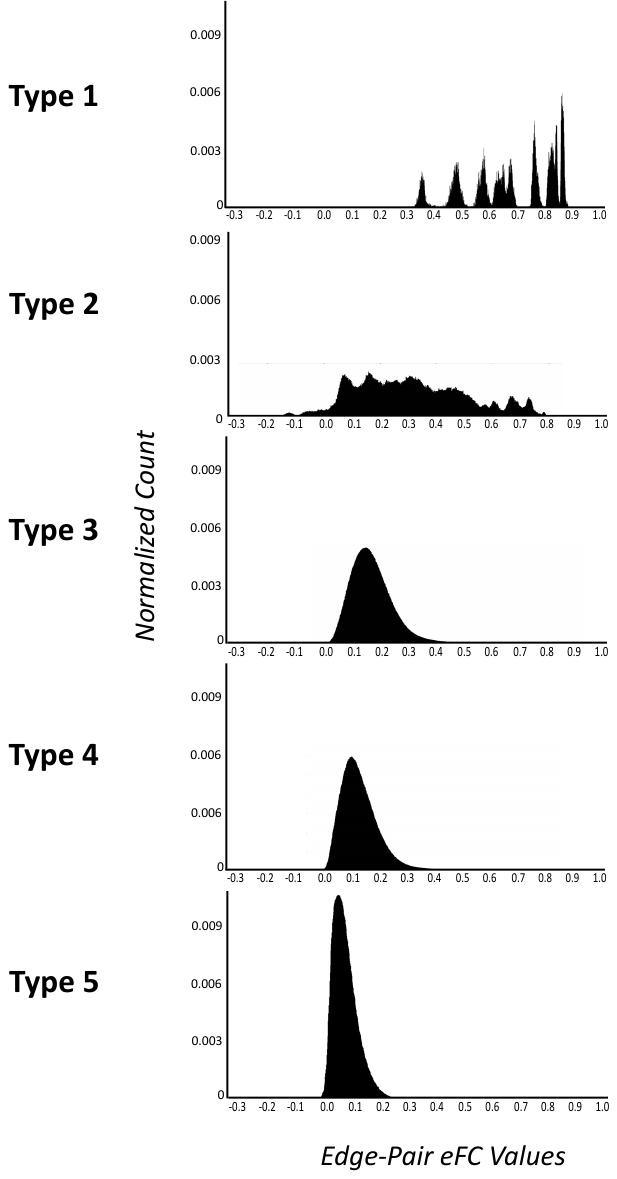

**Figure S3**. Distribution of eFC values for each of the five examined corticostriatal edge-pair types. Each plot displays eFC values (x-axis) separated into 1000 bins. Y-axis represents the normalized (between 0 and 1) count of edge-pairs with eFC values in each range.

Figure S4.

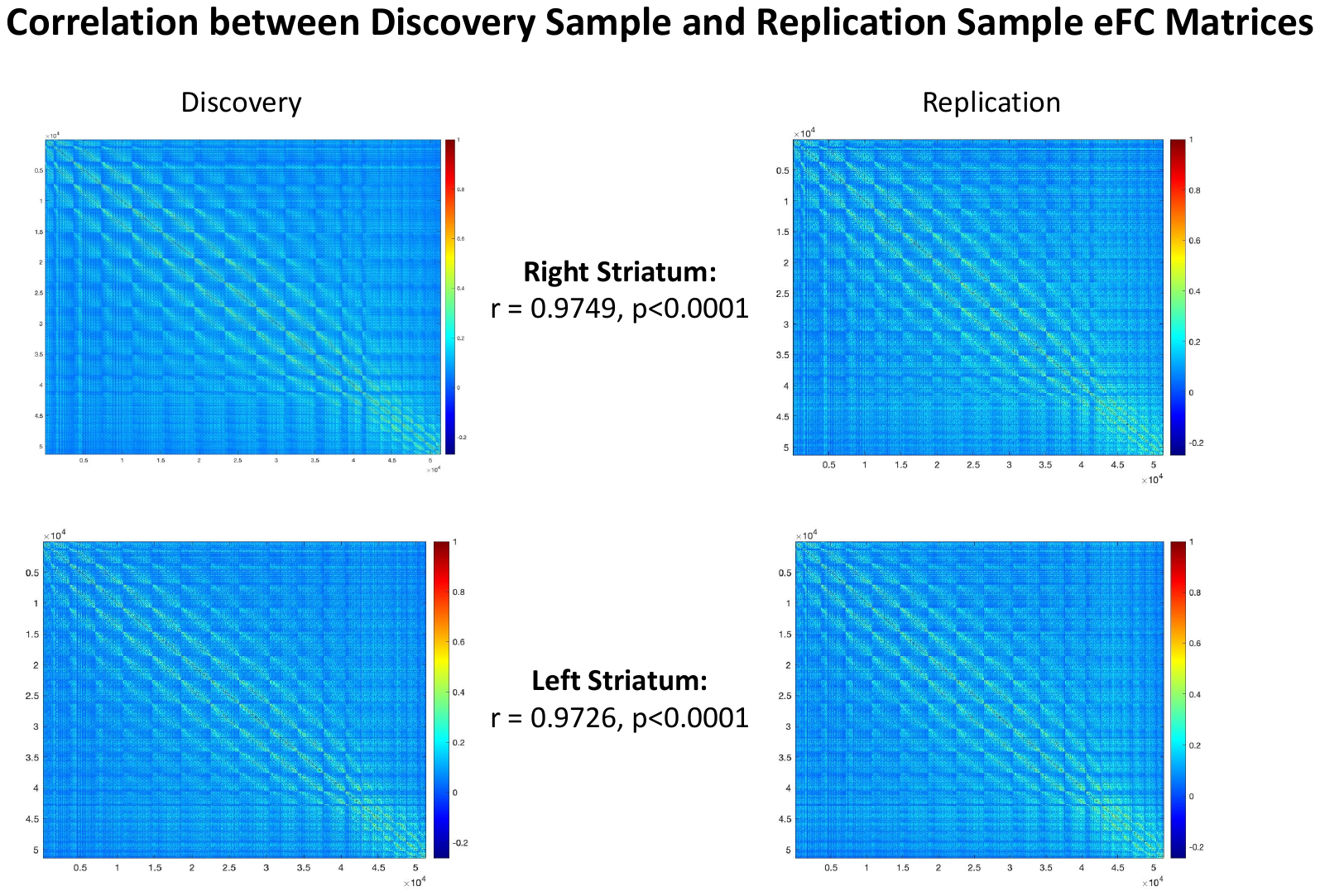

**Figure S4.** Demonstrating the high replicability (as measured by spatial correlation) of corticostriatal temporal organization across distinct, non-overlapping groups of subjects.

Figure S5.

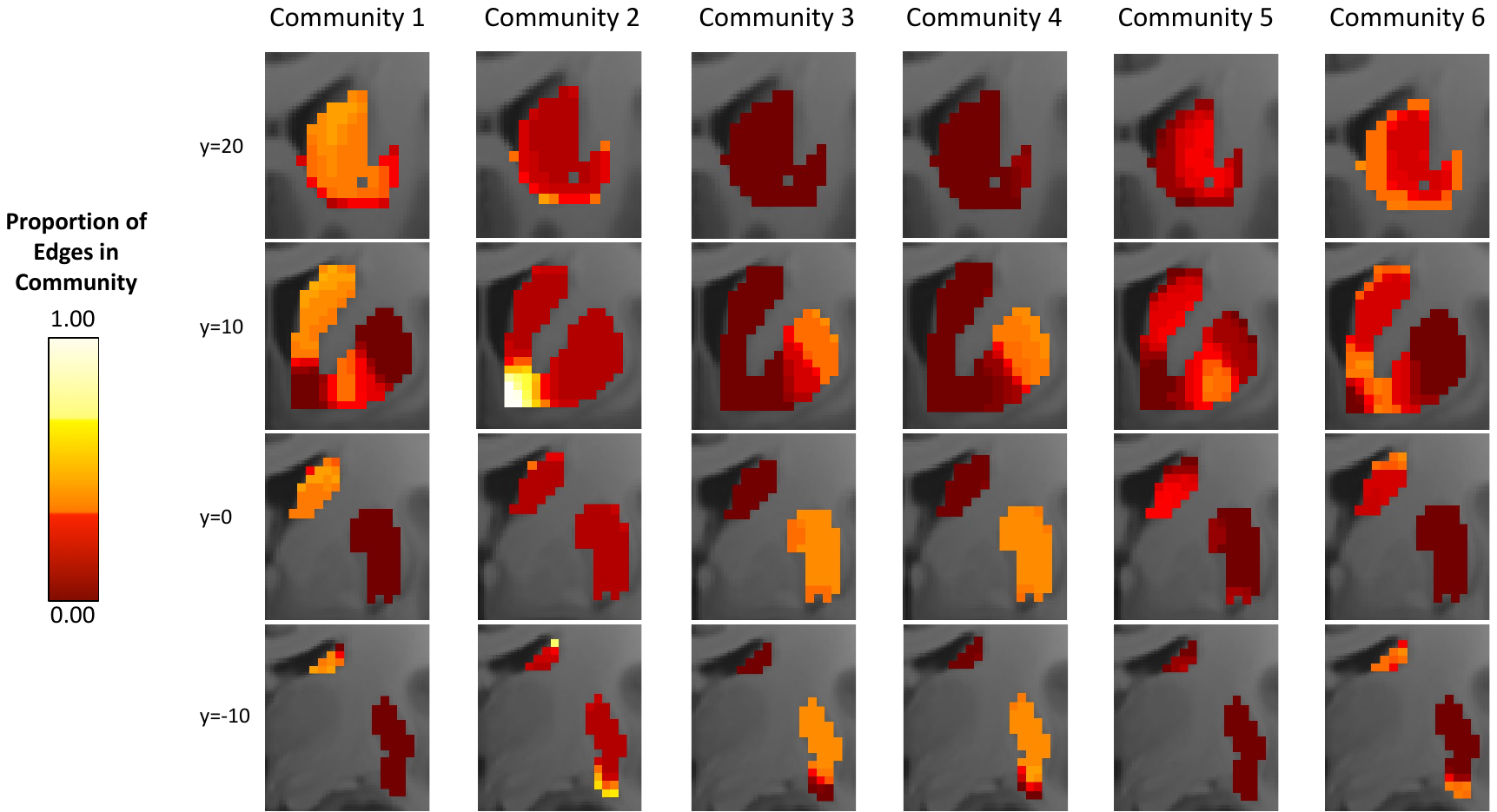

**Figure S5.** Each striatal voxel is a node in 30 different corticostriatal edges. Each edge is assigned, via *k*-means clustering, to one of *k* coactivation synchrony communities. Thus, each striatal voxel can be defined by the proportion of its 30 edges that belong to each community at each level of *k* (*k*=6 depicted here). Increasing brightness indicates a higher proportion of the voxel’s edges belong to that community, and vice versa.

Figure S6.

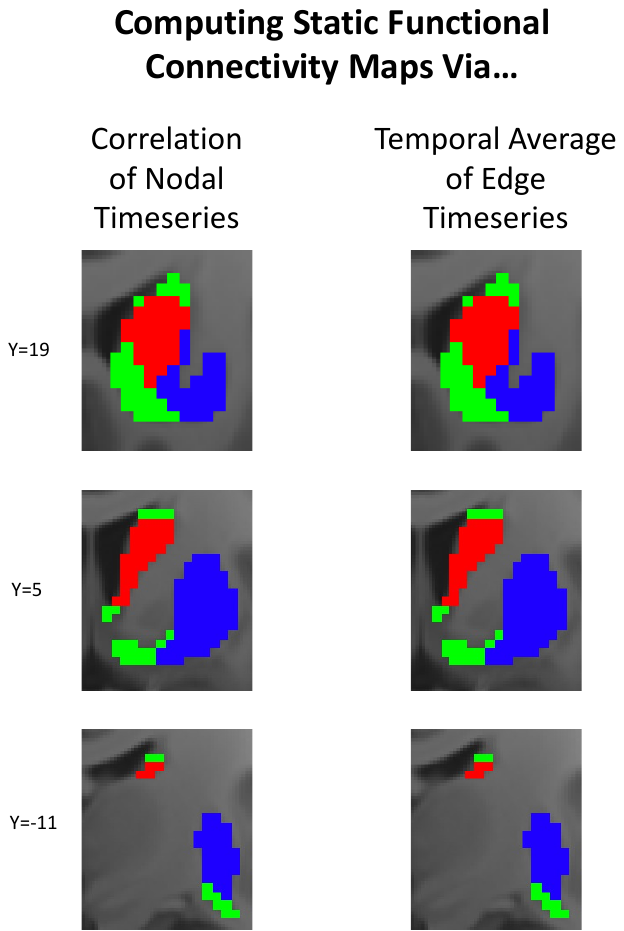

**Figure S6.** The functional connectivity value of a striatal voxel with a cortical ROI can be computed in two ways. In the traditional method, the correlation is computed between the voxel’s BOLD timeseries and the ROI’s timeseries. Here, we show that computing the temporal average of the voxel’s frame-wise coordination magnitudes with the ROI (i.e., the average value of the edge time series) yields precisely equivalent functional connectivity values for each voxel. Thus, both connectivity profile maps and communication pattern maps can be computed from the same input data (i.e., the edge time series data).

Figure S7.

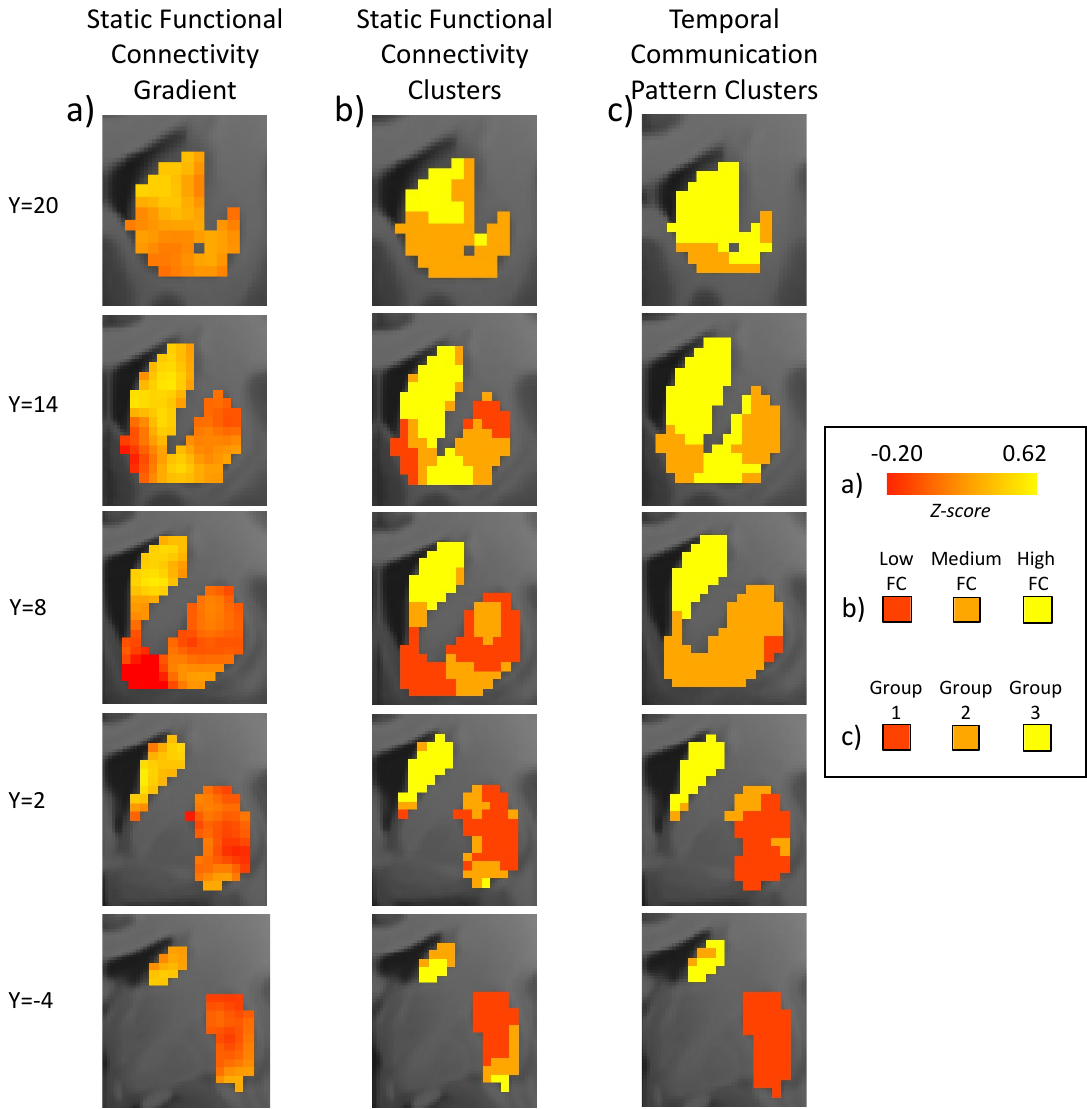

**Figure S7.** Striatal voxels with similar static functional connectivity values with a cortical ROI (a-b) do not necessarily interact with the ROI in the same temporal fashion (c). And vice versa, striatal voxels that interact with a cortical ROI in a similar temporal fashion do not necessarily have similar static functional connectivity values with the ROI. Data shown for a representative subject and ROI (anterior cingulate cortex).

Figure S8.

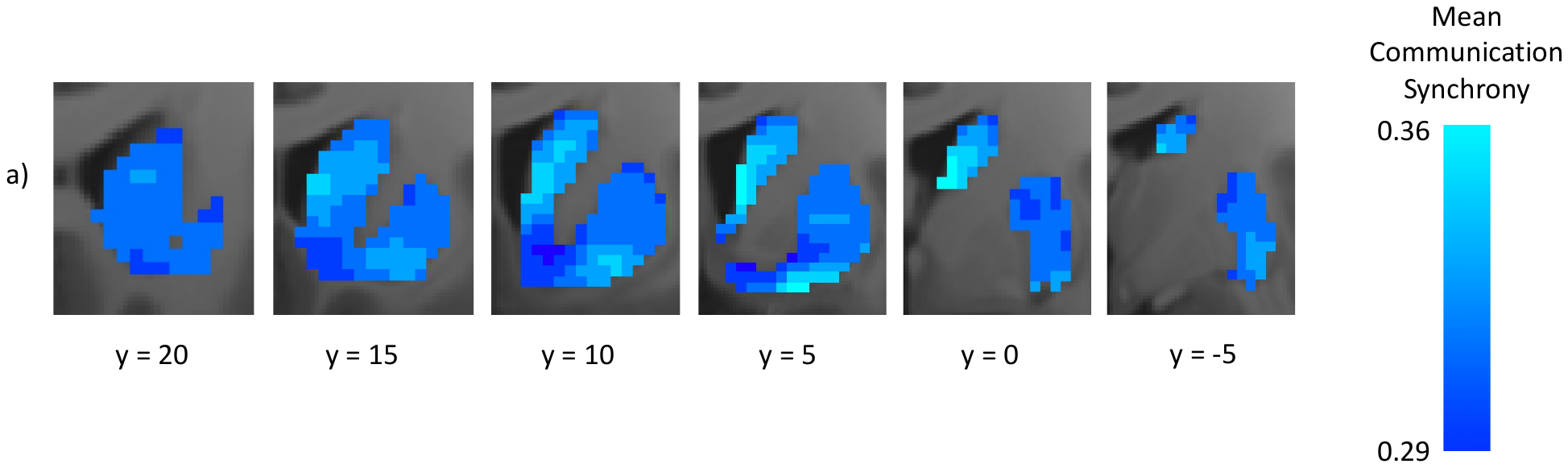

**Figure S8.** The “synchrony” of a striatal voxel’s frontocortical communications was defined as the average eFC amongst all paired combinations of its 30 edge time series. Striatum-wide map of synchrony values, with increasing brightness indicating higher synchrony, and vice versa.
